# sangeranalyseR: simple and interactive analysis of Sanger sequencing data in R

**DOI:** 10.1101/2020.05.18.102459

**Authors:** Kuan-Hao Chao, Kirston Barton, Sarah Palmer, Robert Lanfear

## Abstract

**Summary:** sangeranalyseR is an interactive R/Bioconductor package and two associated Shiny applications designed for analysing Sanger sequencing from data from the **ABIF** file format in R. It allows users to go from loading reads to saving aligned contigs in a few lines of R code. sangeranalyseR provides a wide range of options for a number of commonly-performed actions including read trimming, detecting secondary peaks, viewing chromatograms, and detecting indels using a reference sequence. All parameters can be adjusted interactively either in R or in the associated Shiny applications. sangeranalyseR comes with extensive online documentation, and outputs detailed interactive HTML reports.

**Availability and implementation:** sangeranalyseR is implemented in R and released under an MIT license. It is available for all platforms on Bioconductor (https://bioconductor.org/packages/sangeranalyseR) and on Github (https://github.com/roblanf/sangeranalyseR).

**Supplementary information:** Documentation at https://sangeranalyser.readthedocs.io/.

## 1 Introduction

Sanger sequencing (Sanger and Coulson, 1975; Sanger *et al.*, 1977; Green *et al.*, 2017) was the first controllable method to determine nucleic acid sequences and was commercialized by Applied Biosystems in 1986. Although it has been more than forty years since it was first proposed in 1977, and many new sequencing methods have since been proposed and commercialised, it is still widely-used and indispensable for sequencing individual DNA fragments (Stucky, 2012; Kircher and Kelso, 2010).

Here, we present sangeranalyseR: an automated R package for the analysis of Sanger sequencing data. sangeranalyseR provides: quality trimming, base calling, chromatogram plotting, assembly of contigs from any number of forward and reverse reads, contig alignment, phylogenetic tree reconstruction, and a number of additional methods to analyse reads and contigs in more detail. The package includes two interactive local Shiny apps which allow users to look in detail into each read and contig and change input parameters such as trimming. sangeranalyseR is available on Bioconductor (Gentleman *et al.*, 2004; Huber *et al.*, 2015) (https://bioconductor.org/packages/sangeranalyseR) and is free and open source, which sets it apart from many other packages that perform similar functions, such as Geneious (Kearse *et al.*, 2012), CodonCode Aligner (CodonCode Corcporation, 2003), Phred-Phrap-Consed (Ewing *et al.*, 1998; Ewing and Green, 1998; Gordon *et al.*, 1998), and Sequencher(Gene Codes Corporation, 1991).

sangeranalyseR builds extensively on the excellent sangerseqR (Hill *et al.*, 2014a) package. This package is the only package dedicated to the analysis of Sanger sequencing data on Bioconductor. While sangerseqR is focussed on the analysis of individual Sanger sequencing reads, sangeranalyseR focusses on constructing contigs from multiple Sanger sequencing reads.

## 2 Software description

sangeranalyseR is an R package that can accept input files in **ABIF** or **FASTA** formats. Since the **FASTA** format doesn’t contain the raw data used to call base sequences, when **FASTA** files are used as input some features like trimming, chromatogram plotting, and base calling are not available. R commands in sangeranalyseR can be run on all platforms, and the Shiny applications can be run on all platforms that have a graphical user interface. The package heavily depends on a wide range of R packages from CRAN and Bioconductor (Paradis and Schliep, 2018; Wright, 2016; Wickham, 2007; Schliep, 2011; Schliep *et al.*, 2017; Hill *et al.*, 2014b; Xie *et al.*, 2018; Charif and Lobry, 2007).

sangeranalyseR provides three well-defined S4 classes, *SangerRead, SangerContig* and *SangerAlignment*. These classes form a hierarchy in which a *SangerContig* is defined as a set of one or more *SangerRead*s, and a *SangerAlignment* is a set of two or more *SangerContig*s. User-facing functions in the R package are designed to automate most standard tasks with sensible but adjustable default values. The package has thorough documentation in both a vignette and online using ReadtheDocs. It also includes a large suite of unit tests managed by testthat (Wickham, 2011), and automated by Travis CI (Travis CI Builder Team, 2012). sangeranalyseR is open source, and maintained on Github: https://github.com/roblanf/sangeranalyseR.

## 3 Sequence analysis process

sangeranalyseR needs two kinds of input information: Sanger sequencing reads in either **ABIF** or **FASTA** format, and information on how these reads should be grouped into contigs. The latter information can be provided implicitly with widely-used naming conventions in which the start of each filename defines the contig-group, and the end of each filename defines whether each read is in the forward- or reverse-orientation (e.g. Figure 1A). Alternatively, the same information can be provided in a separate CSV file. Given this information, sangeranalyseR will automatically group reads into different contigs. In Figure 1, we demonstrate a simple workflow. Systematically-named reads are first loaded into R (Figure 1A). Contig assembly and alignment can then be performed with a single command (Figure 1F), and the interactive Shiny application can be launched with another single command (Figure 1F) in order to examine the data and adjust parameters interactively (Figure 1B). The Shiny application includes the read trimming plot (Figure 1D), and the sequencing chromatogram with trimmed region in both 3’ and 5’ ends highlighted (Figure 1E) for each read. Finally, the data can be summarised and output (e.g. Figure 1C) as HTML reports, with the assembled contigs output in **FASTA** format.

**Figure 1.**
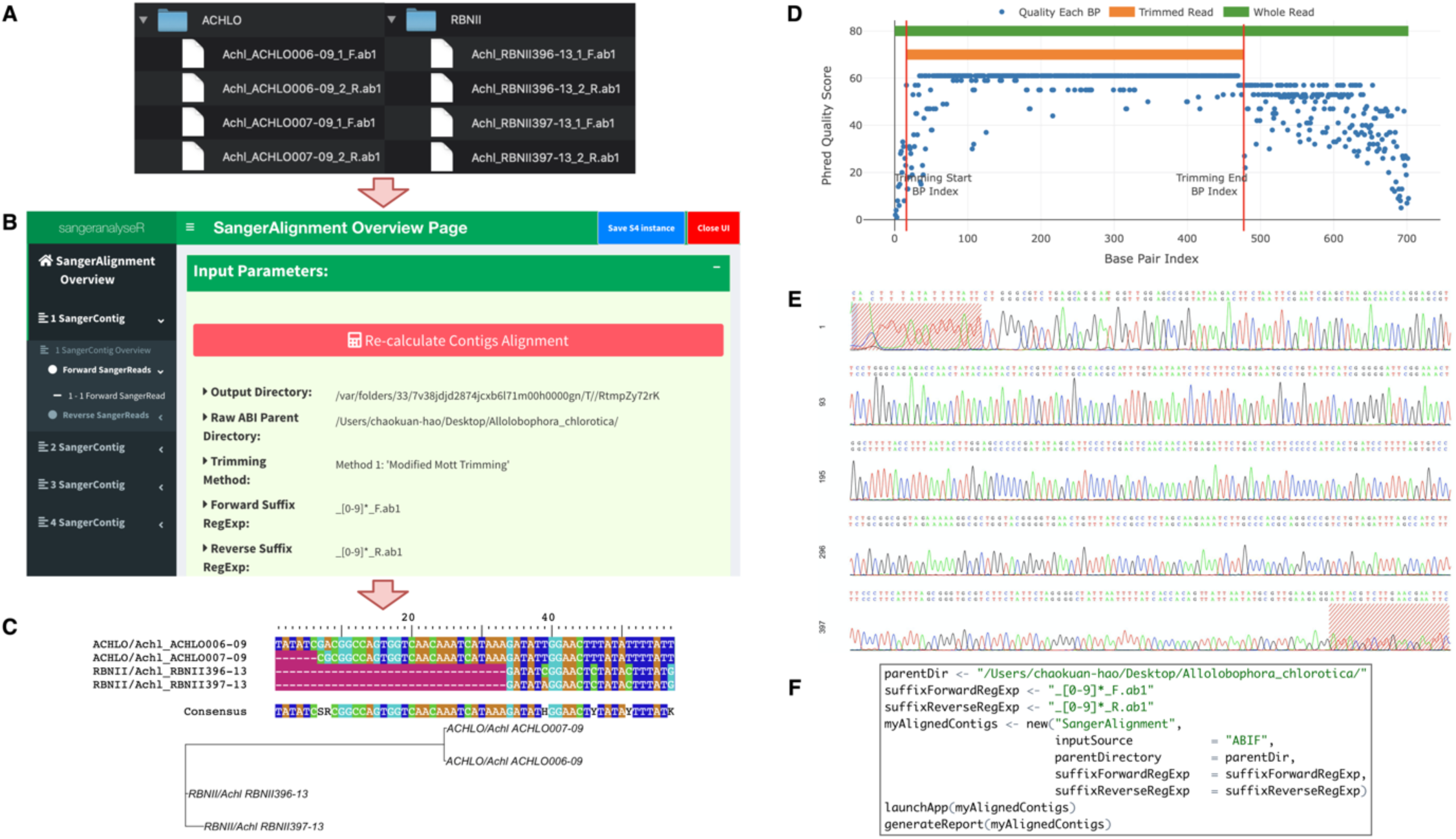
(A), (B), and (C) are SangerAlignment level data analysis workflow. (A) Input files are prepared with a simple file naming convention; (B) The Shiny app allows users to examine each read and each contig, and to adjust trimming and other parameters for each read, using the navigation panel on the left; (C) Each SangerAlignment object provides an alignment of all contigs and a phylogenetic tree of the alignment, to assist users in assessing the quality of the inferences. (D) The read trimming plot shows the Phred Quality score (Y axis) for every base in the read (X axis) along with the trimming locations determined by the trimming parameters (red lines); (E) The chromatogram shows the called bases for each read, as well as the trimmed region at both the 3’ and 5’ ends; (F) An example of the R commands necessary to perform a full analysis, including loading and analysing the reads, launching the Shiny app, and generating the HTML report.

sangeranalyseR includes two methods for trimming low-quality bases from the start and end of each read. The first one is the modified Mott trimming algorithm which is implemented in Phred (Ewing *et al.*, 1998; Ewing and Green, 1998), and BioPython (Cock *et al.*, 2009). The second one is the sliding window trimming approach implemented in Trimmomatic (Bolger *et al.*, 2014). The cutoff value for both algorithms can be adjusted with the “Signal Ratio Cutoff” parameter in either R or Shiny applications.

One limitation of sangeranalyseR is that it does not support editing of individual bases in each read. This is primarily due to the limitations of the R environment. We note though that many other applications support such editing (e.g. Geneious), and that edited reads from those applications can be input to sangeranalyseR using the **FASTA** file input option.

## 4 Conclusion

sangeranalyseR is an open source R package that provides a simple but flexible set of options for analysing Sanger sequencing data in R. It is available on Bioconductor and is free and open source. We hope it will be beneficial to the community and make the analysis of Sanger sequencing simpler and more reproducible.

## Acknowledgements

We thank members of the Palmer and Lanfear labs and a number of beta testers for their feedback on developing the package

